# αB-crystallin affects the morphology of Aβ(1-40) aggregates

**DOI:** 10.1101/2021.03.07.433908

**Authors:** Henrik Müller, David M. Dias, Anna van der Zalm, Andrew J. Baldwin

## Abstract

αB-crystallin (ABC) is a human small heat shock protein that is strongly linked to Alzheimer’s disease (AD). *In vitro*, it can inhibit the aggregation and amyloid formation of a range of proteins including Aβ(1-40), a primary component of AD amyloid plaques. Despite the strong links, the mechanism by which ABC inhibits amyloid formation has remained elusive, in part due to the notorious irreproducibility of aggregation assays involving preparations of Aβ-peptides of native sequence. Here, we present a recombinant expression protocol to produce native Aβ(1-40), devoid of any modifications or exogenous residues, with yields up to 4 mg/L *E. coli*. This material provides highly reproducible aggregation kinetics and, by varying the solution conditions, we obtain either highly ordered amyloid fibrils or more disordered aggregates. Addition of ABC slows the aggregation of Aβ(1-40), and interferes specifically with the formation of ordered amyloid fibrils, favouring instead the more disordered aggregates. Solution-state NMR spectroscopy reveals that the interaction of ABC with Aβ(1-40) depends on the specific aggregate morphology. These results provide mechanistic insight into how ABC inhibits the formation of amyloid fibrils.

**Highlights:**

- Protocol for production of native recombinant Aβ(1-40)
- Amyloid formation under physiological conditions is highly reproducible
- Both ordered fibrils and disordered aggregates can be reliably formed
- αB-crystallin specifically inhibits amyloid fibril assembling, favouring disordered aggregates

**eTOC blurb:** Müller *et al.* introduce a protocol for the highly reproducible production of amyloid from native Aβ(1-40) and determine that the human chaperone ABC specifically destabilises them in favour of disordered aggregates. NMR shows that ABC can distinguish between aggregate morphologies.

**Graphical Abstract:** 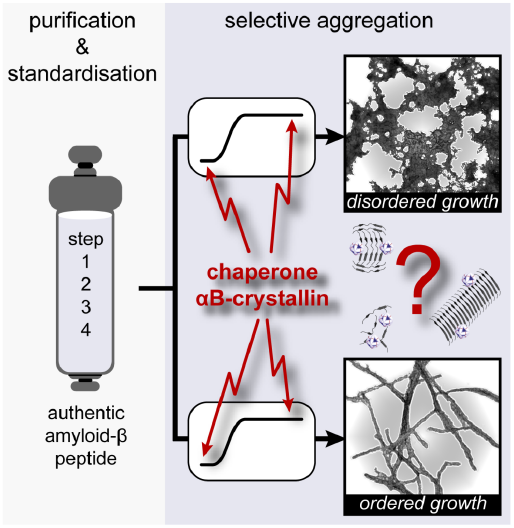

## Introduction

Alzheimer’s disease (AD) is associated with the aggregation of amyloid-β (Aβ)-peptides into amyloid plaques in the brains of AD patients **(Thal, et al., 2015)**. The human small heat shock protein αB-crystallin (ABC) is up-regulated in neurons and glia adjacent to amyloid plaques **(Haslbeck, et al., 2005)** and is co-localised with amyloid plaques extracted from AD patients **(Ecroyd and Carver, 2009)**. *In-vitro*, ABC prevents a range of proteins from aggregating and forming amyloid and reduces the toxic effects of amyloid on cell cultures **(Hochberg, et al., 2014; Wilhelmus, et al., 2006)**. Despite these links, studying its mechanism has proven tremendously challenging. The principle difficulty is in obtaining reproducible kinetic data. Even when isolated, ABC spontaneously adopts a polydisperse range of oligomers containing ca. 10 to 50 subunits centred on a mass of ~ 560 kDa **(Hilton, et al., 2013)**. When mixed with solutions containing aggregates for mechanistic study, the range of populated oligomeric complexes becomes substantially larger, confounding experimental analysis. Given the complexity of isolated ABC, to understand its links to AD, it is crucial to study its effects on physiologically relevant Aβ peptides, under conditions that give rise to reproducible aggregation kinetics.

Aβ-peptides with 36-43 amino acid residues have been identified in the brain and cerebrospinal fluid of AD-patients, of which the most abundant is Aβ(1-40). Studies of Aβ are usually focused on the two peptides with the highest population in plaques, Aβ(1-40) and Aβ(1-42). These are typically prepared by solid-phase synthesis. The specific morphologies of amyloids produced from these two forms and their cytotoxic responses differ considerably (**Bitan, et al., 2003; Pauwels, et al., 2012)**. Moreover, experiments following aggregation kinetics can vary widely between different preparations of source material. This is thought to be due to a combination of factors including impurities, variable oxidation states of Aβ’s internal methionine, the polymorphic nature of the aggregation process, and having variable populations of oligomeric Aβ-aggregates when the reaction is initiated **(Finder, et al., 2010; Hou, et al., 2004; Zagorski, et al., 1999**). Studying the effects of mutations, variations in solution conditions, and binding partners remains challenging **(Luhrs, et al., 2005; Yonemoto, et al., 2009)**. Data including kinetic aggregation curves, cyto-toxicities and models of fibril architecture can differ considerably between laboratories and preparation method **(Cohen, et al., 2015; Fandrich, et al., 2011; Luhrs, et al., 2005; Mainz, et al., 2015; Miller, et al., 2010; Paravastu, et al., 2008; Ravotti, et al., 2016; Tycko, et al., 2009; Xiao, et al., 2015)**, and variations between amyloid fibrils produced *in vivo*, and those derived from brain tissue samples from AD patients have been observed **(Paravastu, et al., 2009)**.

It is likely that at least in part, variations in results derive from the use of synthesised Aβ-peptides for analysis from different sources. The inadvertent inclusion of small populations of truncations inherent in synthesising a peptide might be expected to add a relatively random element to aggregation kinetics (e.g. **Fig. 2**). To overcome this limitation, protocols have been developed to produce Aβ-peptides recombinantly. These methods seek to purify Aβ either from inclusion bodies, in which Aβ-peptides have an additional N-terminal methionine **(Cohen, et al., 2015; Walsh, et al., 2009)** or by fusion to a solubilising protein domain and/or polyhistidine tag **(Finder, et al., 2010; Garai, et al., 2009; Hortschansky, et al., 2005; Long, et al., 2011; Weber, et al., 2014; Wiesehan, et al., 2007; Zhang, et al., 2009)**. These methods can lead to non-physiological sequences because of the cleavage enzyme used **(Garai, et al., 2009; Long, et al., 2011; Wiesehan, et al., 2007; Zhang, et al., 2009)**, or the resulting material can show variation in the aggregation kinetics **(Finder, et al., 2010; Hortschansky, et al., 2005; Weber, et al., 2014).** Recombinant Aβ-peptides can lead to highly reproducible kinetic behaviour, and the methionine protocol has been analysed in detail to provide a detailed mechanistic characterisation of amyloid formation. Notably, the mechanism by which Aβ(1-42) assembles into amyloids follows secondary nucleation kinetics, where the surfaces of amyloids effectively act as seeds to accelerate amyloid formation **(Cohen, et al., 2015; Cohen, et al., 2013)**.

Here, we provide an inexpensive protocol to produce a recombinant source of pure and entirely native Aβ(1-40) without any exogenous amino acid residues or modifications in relatively high yield. Aβ(1-40) is expressed in the soluble fraction as a fusion protein with a small ubiquitin-like modifier (SUMO) domain and is cleaved to give exactly the native sequence. We demonstrate that it can be induced to reliably form a range of aggregate morphologies that depend on precisely how the aggregation reaction was initiated. Either straight discrete amyloids fibrils can be formed, which we term here type 1 (**Fig. 3**), or less ordered non-uniform aggregates, a morphology that we term type 2 (**Fig. 4**). Notably, the kinetics of assembly of both aggregate types is shown to be highly reproducible. The material allows us to study the interactions of Aβ(1-40) aggregates with ABC using both fluorescence and solution-state NMR, revealing that the ABC interaction is sensitive to the specific morphology of Aβ-aggregates. We demonstrate that ABC selectively blocks the formation of amyloid fibrils, leading Aβ to preferentially form more disordered aggregates. We present a model for the Aβ and ABC interaction that is consistent with both our observations here, and recent mechanistic studies **(Cohen, et al., 2013; Mainz, et al., 2015; Narayanan, et al., 2006) (Fig. 7)**, where Aβ(1-40) and ABC are effectively in competition for terminal binding sites, and lateral surface sites on aggregates.

## Results

### Purification of native Aβ(1-40)

To prepare highly pure, homogeneous, native Aβ (1-40), we fused the small ubiquitin-like modifier (SUMO) to the N-terminus of the Aβ(1-40) peptide (**Fig 1A**). The SUMO protease Ulp1 recognises the tertiary structure of SUMO and, in turn, cleaves at an exactly prescribed position (**Malakhov, et al., 2004**). SUMO has been harnessed to provide a robust method to enhance the solubility and stability of recombinant proteins **(Satakarni and Curtis, 2011; Yates, et al., 2016)**. Further, a double poly-histidine tag was placed at the N-terminus of SUMO, resulting in His-SUMO-Aβ, which facilitates purification **(Fischer, et al., 2011; Khan, et al., 2006)**. To illustrate its versatility, we developed two methods for purification of Aβ(1-40) (experimental procedures), one based on the denaturant GdnHCl (**Fig 2**) and one based primarily on the Tris buffer (**Fig S1**). The yield was similar in both cases, up to 4 mg/L *E. coli,* and the aggregation behaviour of material from both methods was found to be highly similar, and so we describe both here. The protocols require only standard chromatographic equipment, are amenable to upscaling, and can be adapted to generate a range of Aβ peptides.

**Fig. 1:**
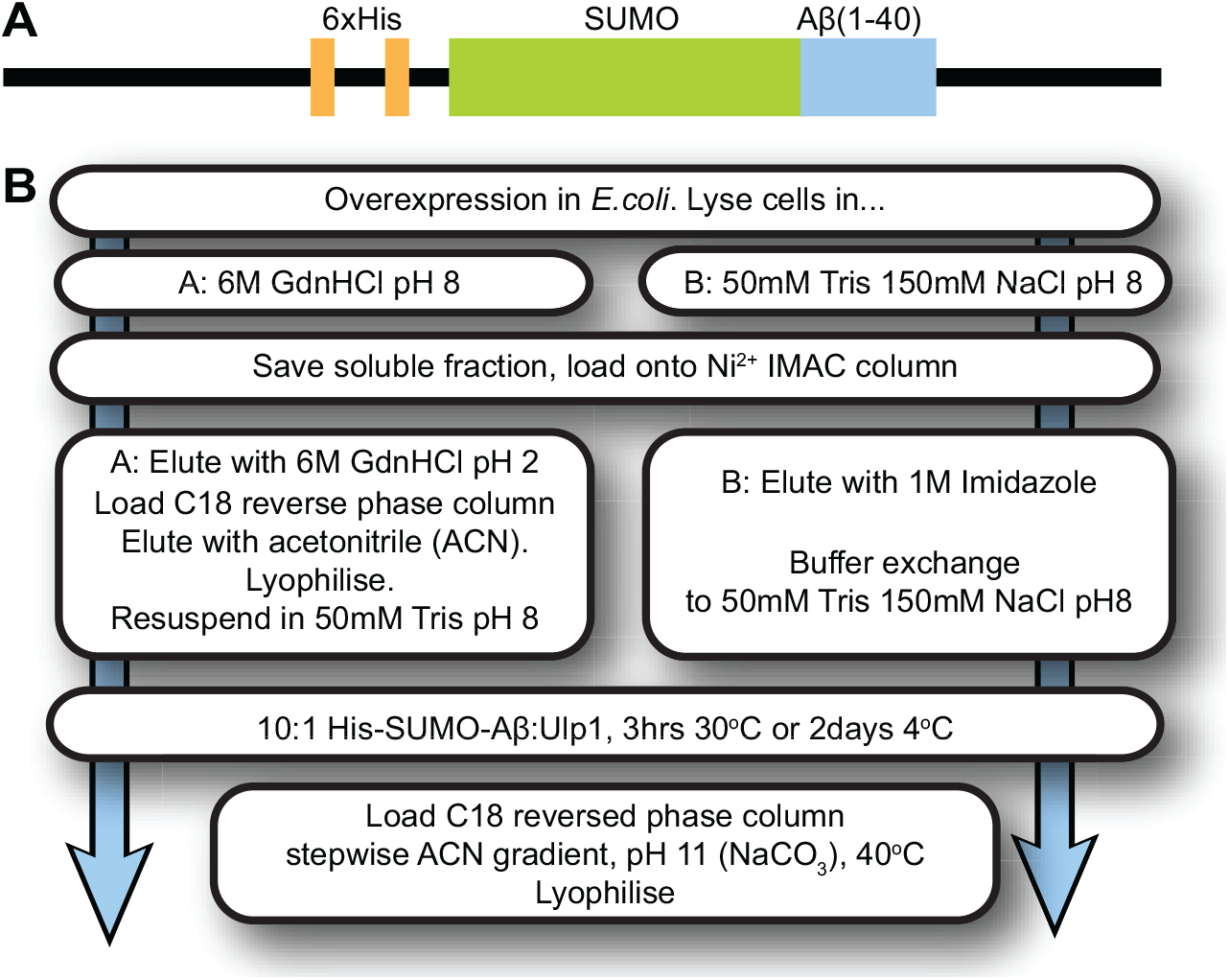
Expression His-SUMO-Aβ(1-40) inside pET-28a(+) and workflow for purification of Aβ(1-40). **(A)** Schematic representation of a linearised pET-28a(+) vector containing two hexahistidine - affinity tags, an 11-amino acid residue linker, a SUMO tag, and full-length Aβ(1-40). **(B)** Flowchart of key steps of purification of Aβ(1-40). Two purification methods are described, one using GdnHCl solutions (**Fig 2**), and one using Tris buffers (**Fig S1**). The yields of both were found to be similar (2-4 mg / L *E. coli*) and no difference in aggregation behaviour was observed, and so we describe both in this article. The final Aβ(1-40) sequence DAEFRHDSGYEVHHQKLVFFAEDVGSNKGAIIGL MVGGVV is exact, as the protease we use to remove the tag cleaves precisely where desired and the starting methionine is removed with the N-terminal tag. The aggregation kinetics of material from both methods was identical. The molecular weight of monomeric Aβ(1-40) of 4329.1 ± 2.4 Da, as measured by ESI-MS, is identical to the expected 4329.8 Da (**Fig S2**).

**Fig. 2:**
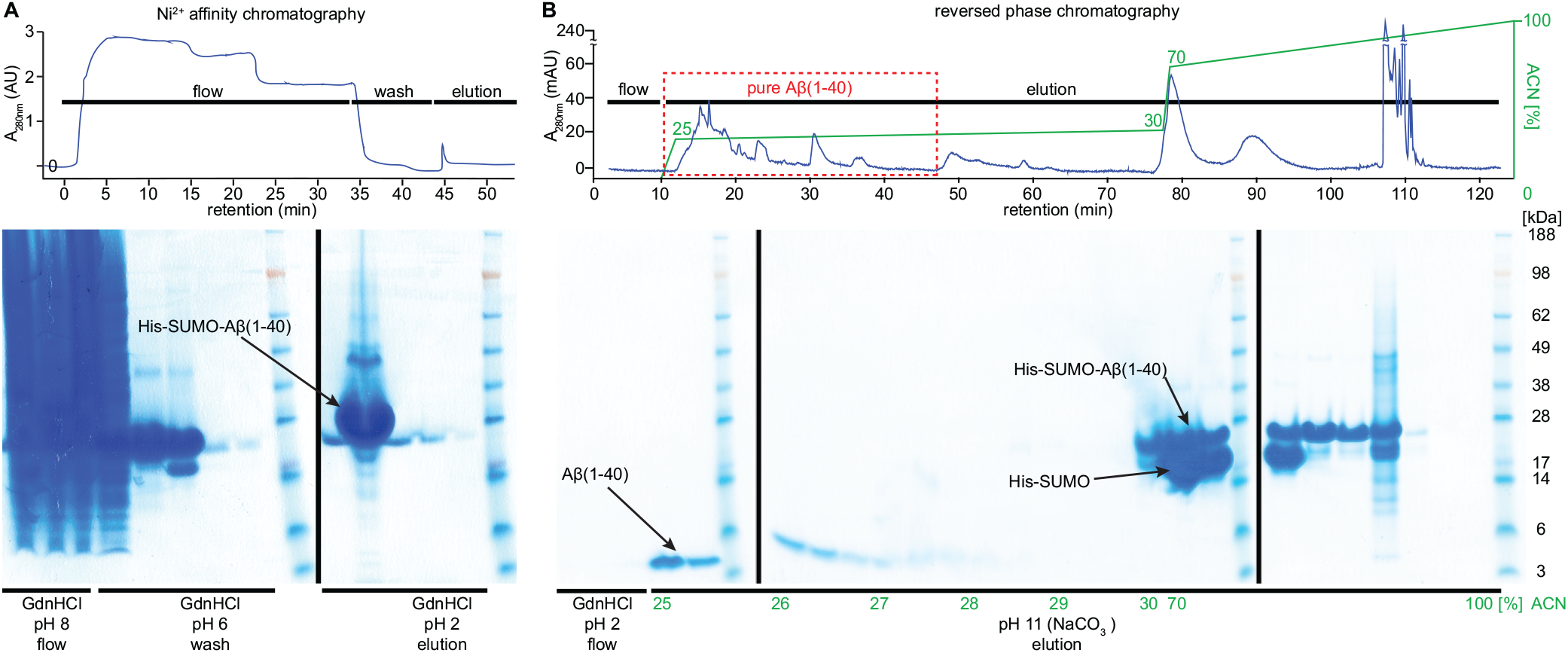
The two chromatography steps for purification of Aβ(1-40). **(A)** Typical performance of purification steps using the GdnHCl method (**Fig 1B, methods**). The cell lysate from 1 L *E. coli* growth was loaded onto a 5ml HisTrap column containing immobilised Ni^2+^. After washing with 6 M GdnHCl pH 6, His-SUMO-Aβ was eluted with 6 M GdnHCl pH 2. The buffer was exchanged to 50mM Tris pH 8 for cleavage of the SUMO-Aβ linker with Ulp1 for 3 hours at 30 °C. **(B)** After digestion, the reaction was diluted into 6 M GdnHCl pH 2 and separation of His-SUMO-Aβ from Aβ was achieved by reversed phase chromatography. A stepwise gradient from 25 - 30 % (vol/vol) acetonitrile at in 10 mM NaCO_3_ at pH 11 and 40 °C was applied to elute purified Aβ(1-40).

In brief, *E. coli* cells expressing His-SUMO-Aβ are lysed and the soluble fraction is loaded on a column containing immobilised Ni^2+^ to attract the His tag (**Fig. 2A** and **Fig. S1A**). Upon elution, optimal cleavage of His-SUMO-Aβ was achieved by adding Ulp1 at a molar ratio of 10:1 for either 30 °C for 3 h or at 4 °C for 2 days. We did not observe the addition of detergents, altered salt concentrations or altered pH to improve the yield of cleavage.

Separating His-SUMO-Aβ from cleaved Aβ was surprisingly challenging, likely owing to the exceptional tendency of Aβ to self-assemble. We were able to obtain highly pure Aβ(1-40) only after binding to a C18 reversed phase column and eluting with a stepwise acetonitrile (ACN) gradient ranging from 25 - 30 %, whereas His-SUMO-Aβ eluted at ACN concentrations higher than 30 %. This separation depended crucially on a high pH of 11 and was further enhanced by holding the column at an elevated temperature of 40 ^o^C (**Fig. 2B** and **Fig. S1B**).

SDS-PAGE of final samples demonstrated the correct migration of a single band at 4.3 kDa. Both staining with InstantBlue, and densitometric analysis confirmed that the final Aβ(1-40) was more than 97 % pure. This finding was corroborated by electrospray ionisation mass spectrometry confirming one final product with a molecular weight of 4329.1 ± 2.4 Da, essentially identical to the one expected for native Aβ(1-40) of 4329.8Da (**Fig. S2**).

### Robustness and consistency of aggregation kinetics

To assess the aggregation kinetics of purified recombinant Aβ(1-40), we combined light scattering measurements with a continuous thioflavin T (ThT) binding assay, a dye whose fluorescence is enhanced on binding amyloid **(LeVine, 1999; Naiki, et al., 1989)**, and negative-stain transmission electron microscopy (TEM, **Fig. 3**). Aggregation was initially triggered by dissolving lyophilised peptide in 10 mM sodium phosphate pH 7.4 and 100 mM sodium chloride, taking to pH 11 with 10 mM NaOH, holding at 4 °C for 30 min, returning to pH 7.4 by the addition of 10mM HCl, and incubating at 37 °C at 700 rpm linear shaking **(Ryan, et al., 2013)**, a protocol that promotes the formation of homogeneous amyloid fibrils (**Fig. 3**). Under these conditions, we obtained kinetic traces typical for protein aggregation. A lag phase was first observed, during which any present aggregates were below the detection limit of the optical equipment, followed by a rapid increase in signal intensity and a final stationary phase. We typically observe a slight decrease in ThT fluorescence intensity at longer times, which we can tentatively attribute to lateral fibril association resulting blockage of ThT binding sites. Only homogeneous, discrete amyloid fibrils were observed in TEM micrographs (**Fig. 3**).

**Fig. 3:**
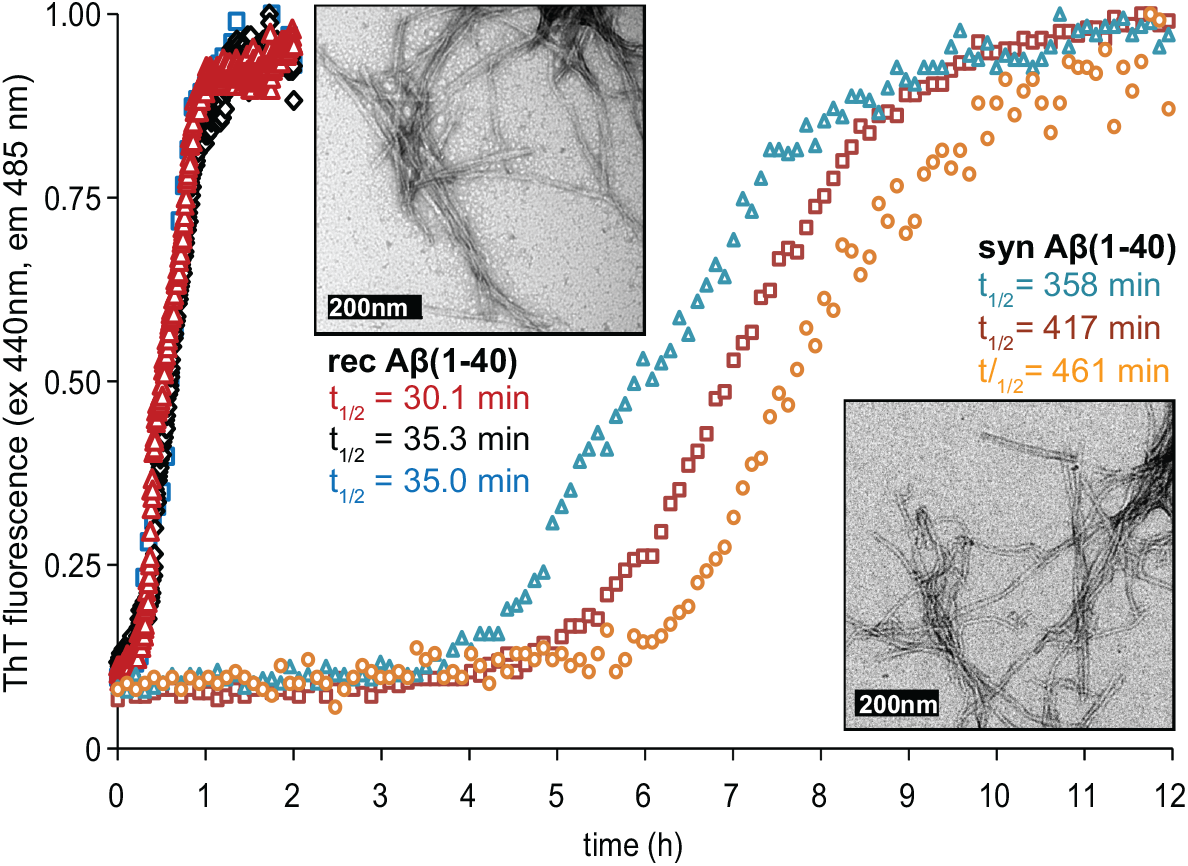
Recombinant Aβ(1-40) has highly reproducible aggregation kinetics. Three samples of 20 μM recombinant Aβ(1-40) were produced at different times in a 4-month period, and treated with 10 mM NaOH at 4 °C for 30 minutes following a previously established protocol for homogeneous amyloid fibrils **(Ryan, *et al.*, 2013)**, before neutralisation and aggregation in 10 mM NaP_i_, 100 mM NaCl pH 7.4. The aggregation reactions were performed up to 4 months apart at 37 °C and 700 rpm linear shaking. No smoothing, fitting, or averaging procedures was applied to the data shown. The aggregation curves of the recombinant Aβ were highly similar. By contrast, three samples of 20 μM synthetic Aβ(1-40) from a single batch were exposed to identical conditions. The respective times to half completion are indicated. TEM micrographs were prepared by applying 5 μl of the mixtures to the EM grids once the reaction had reached a steady-state. Amyloid fibrils from both were identical by visual inspection of TEM micrographs, although the aggregation kinetics from the synthetic peptides showed significant variation in repeats as shown.

To quantify the reproducibility of our protocol, we acquired three datasets recorded from independent samples prepared months apart (**Fig. 3**). These traces were highly similar. We determined the times to half completion for each individual data set by extracting the data points corresponding to a ThT-fluorescence value between 0.3 and 0.7 and performing a linear regression **(Cohen, et al., 2013)**. The average half completion time was 33.5 ± 2.4 min. We tested a commercially available synthetic Aβ(1-40) from a single batch and exposing it to identical solution conditions in triplicate, resulting in a half completion time of 412 ± 42 min (**Fig. 3**). The decrease in the total length of the half completion time by a factor of 12 and the higher consistency demonstrates the reliability of our Aβ(1-40) purification protocol. TEM micrographs of both samples showed amyloid fibrils, which we could not distinguish qualitatively, although structural differences may exist **(Paravastu, et al., 2009)**.

For comparison, half completion times from aggregation assays conducted under similar conditions ranged between 1 and 24 hours and can vary from batch to batch by up to several hours **(Ecroyd and Carver, 2009; Hortschansky, et al., 2005; Mainz, et al., 2015; Shammas, et al., 2011)**. The highly reproducible aggregation kinetics and native sequence from Aβ(1-40) produced by our protocol renders it highly amenable for mechanistic analysis.

### Producing aggregates with specific morphologies

We acquired kinetic data for aggregation, varying Aβ(1-40) concentrations at pH 7.4 and 37 °C performed either with or without a prior 30 min at 4 °C incubation at pH 11. A combination of solution scattering measurements, ThT fluorescence, and TEM were used to characterise the aggregation progress. Two distinct types of aggregates (type 1 and type 2) were observed depending on the solution conditions used. Type 1 contained only straight discrete amyloid fibrils of a width ca. 10 nm and lengths of up to several μm (**Fig. 4A**). These type 1 aggregates are characterised by a relatively high ThT fluorescence and relatively low light scattering. By contrast, type 2 aggregates show a more disordered non-fibrillar and non-uniform morphology in TEM micrographs, relatively little ThT binding, and pronounced light scattering (**Fig. 4B**). We attribute these differences to the reduced scattering of rod-like (type 1) over spherical (type 2) particles and the higher surface area-to-volume of rod-like molecules for ThT binding (type 1) over the spherical (type 2) aggregates. Moreover, the morphology of aggregates was highly reproducible, depending on the solution conditions. Pure type 1, pure type 2, and mixtures of the two were produced under well-defined purification conditions (see discussion).

**Fig. 4:**
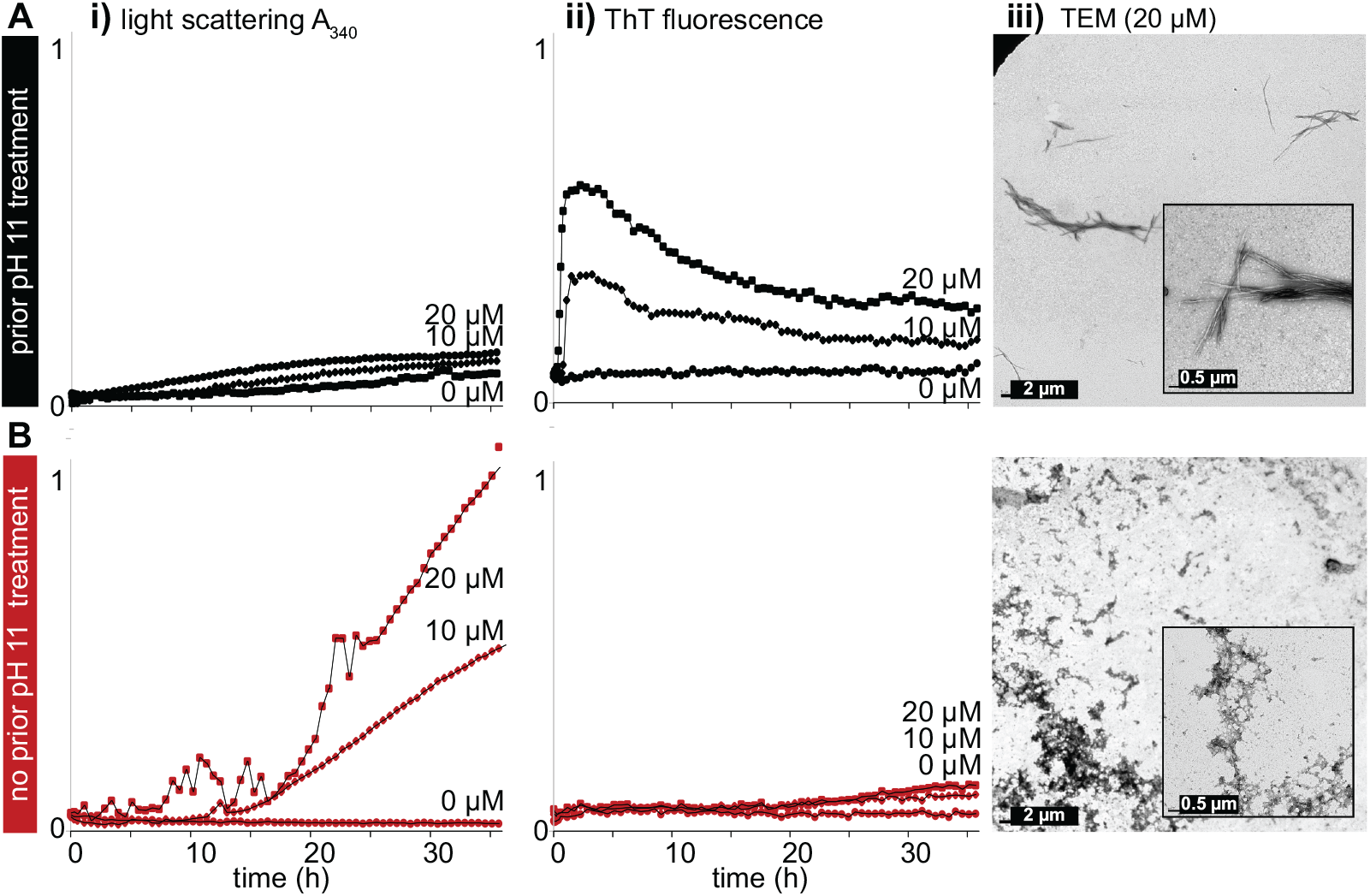
Initial pH treatment significantly affects the morphology of Aβ(1-40) aggregates. Aggregation was monitored simultaneously by light scattering at 340 nm (i) and Th-T fluorescence (ii), and TEM micrographs were acquired once the reaction reached a steady-state (iii). Note that all scattering and florescence curves in the paper (**Fig 4,5 S3, S4**) are normalised in the same way and can be directly compared. The concentration of Aβ(1-40) was varied as specified. Reactions were initiated either with (**A**) or without (**B**) a prior pH treatment, where samples were taken to pH 11 at 4 °C for 30 minutes, following a previously established protocol **(Ryan, *et al.*, 2013)**. The kinetic traces follow aggregation in 10 mM NaP_i_, 100 mM NaCl pH 7.4 at 37 °C. With a prior pH treatment, scattering was relatively low, Th-T fluorescence was relatively high, and discrete amyloid fibrils were observed by TEM. We define this morphology as ‘type 1’. By contrast, without a pH treatment, significantly greater scattering was observed with lower Th-T fluorescence, and non-uniform disordered aggregates were observed by TEM. We define this morphology as ‘type 2’. All reactions were performed in triplicate but only one trace is shown in this figure for clarity (see **Fig. 3** and specific error bars in **Fig. 5**).

In absence of a prior pH 11 treatment, prepared under solution conditions that are physiologically mimicking, we observed only type 2 aggregates (**Fig. 4B**) and no evidence of mature amyloid fibrils (type 1). It is interesting to note that ThT-binding is not restricted to discrete linear amyloid fibrils but has also been observed for oligomeric intermediates (**Carrotta, et al., 2001; Groenning, et al., 2007; Maezawa, et al., 2008**). As we observed ThT fluorescence from these morphologically non-fibrillar type 2 aggregates, they have features that are characteristic of amyloid (**Fig. 4Bii**).

By contrast, when we subjected ≤ 20 μM Aβ(1-40) to a treatment at pH 11 prior to aggregation, we observed only type 1 aggregates, i.e. discrete amyloid fibrils, in TEM micrographs (**Fig. 4A**). At concentrations ≥ 20 μM Aβ(1-40), a mixture of both morphologies was detected (**Fig. S3**). Taken together, we conclude that our recombinant Aβ(1-40) can be used to produce reproducibly either two distinctive types of aggregates or mixtures of the two.

To further investigate how generation of either amyloid fibrils or non-fibrillar aggregates depended on the chosen buffer conditions, we repeated our kinetic analysis using 50 mM NH_4_-acetate pH 8.5, instead of 10 mM NaP_i_ 100 mM NaCl pH 7.4 (**Fig. S4**). Despite the presence of different salts, different salt contents, and different pH-values, the overall trends were highly similar. We always observed pure discrete type 1 amyloid fibrils below 20 μM Aβ(1-40) after a prior pH treatment, a combination of both aggregate types at higher Aβ(1-40) concentrations, and only non-fibrillar type 2 aggregates in the absence of a prior high pH treatment.

### Chaperone ABC can distinguish between both types of aggregate

Having established conditions to produce distinctive types of Aβ aggregates reliably and reproducibly, we sought to determine effects of ABC on their aggregation. We incubated Aβ(1-40) with ABC and, as has been reported previously **(Ecroyd and Carver, 2009; Hochberg, et al., 2014)**, observed a considerable delay in the onset of aggregation. At longer timescales, however, aggregation was still observed under all conditions tested. In the absence of ABC, both Aβ(1-40) aggregate morphologies were found in the pellet fractions after differential centrifugation owing to their high molecular weight (**Fig. S5A**). By contrast, when ABC was present at the beginning of the aggregation reaction, the vast majority of Aβ(1-40) was relocated to the supernatants indicating a considerable reduction of high molecular weight species. When we investigated the aggregates using TEM, we observed that all aggregates in the presence of ABC were of type 2. Most notably, even under conditions that favour type 1 discrete amyloid fibrils in absence of ABC, its addition at the start of aggregation always led to non-uniform disordered aggregates of type 2 morphology (**Fig. 5**).

**Fig. 5:**
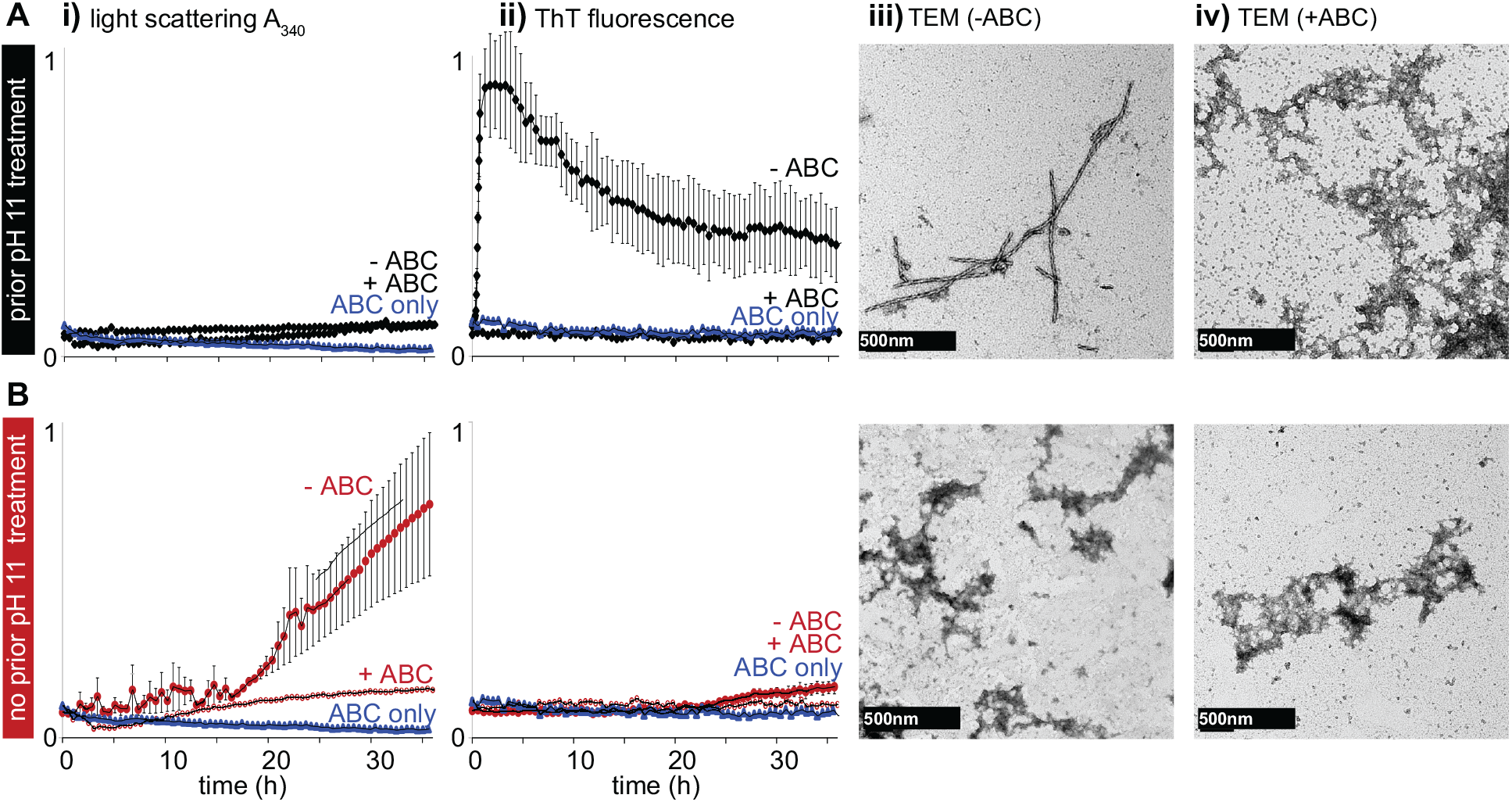
The chaperone ABC slows aggregation, and can affect the morphology of aggregates. Aggregation reactions with (**A**) and without (**B**) a prior pH 11 treatment were followed using light scattering (i), fluorescence (ii) and TEM (iii) as in Fig. 4 in the presence and absence of ABC. Note that all scattering and florescence curves in the paper (**Fig 4, 5, S3, S4**) are normalised in the same way and can be directly compared. In absence of ABC, 20 μM Aβ(1-40) can form either fibrillar type 1, or type 2 amyloids depending on whether or not prior pH 11 treatment was applied (Fig 4). If 5 μM ABC was pre-incubated with either, a significant reduction in both light scattering and ThT fluorescence was observed indicating ABC function as a chaperone. Notably however, in the case of type 1 fibrillar aggregation, fibrils were not observed to appear when samples were pre-incubated with ABC. Instead, aggregates that were morphologically highly similar to type 2 were observed. Light scattering (i) and ThT fluorescence data (ii) were acquired in triplicate and represented as mean ± SEM for each time point.

To study whether ABC has a distinguishable influence on mature Aβ(1-40)-aggregates, we incubated pre-formed type 1 and type 2 aggregates for two days in the presence of ABC. Under these conditions, ABC has been observed to bind to the surface of Aβ fibrils **(Narayanan, et al., 2006; Shammas, et al., 2011)**. Consistent with this, we observed no change in light scattering or ThT-fluorescence or the morphology of Aβ-aggregates (**Fig. 6**).

**Fig. 6:**
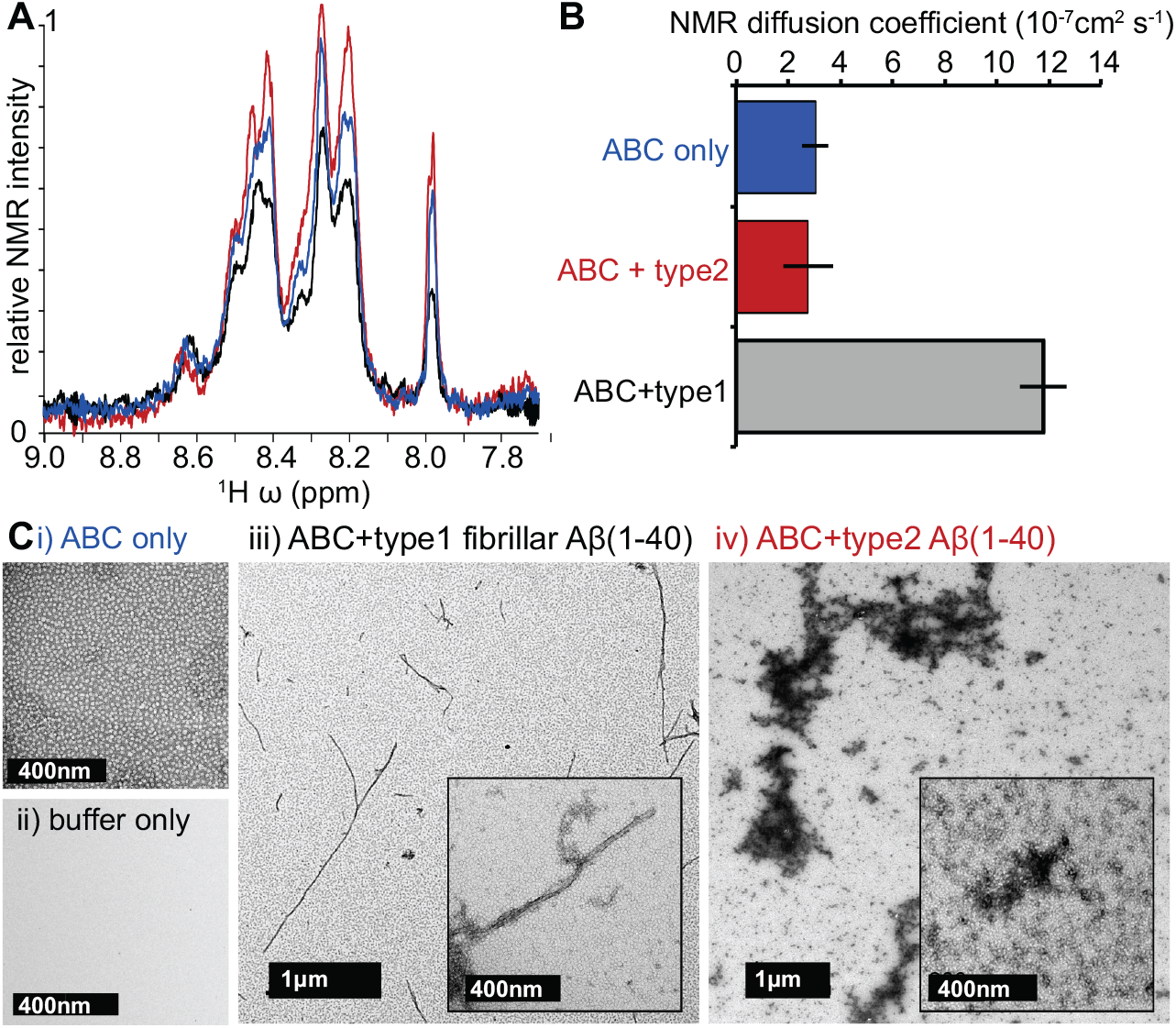
ABC interactions vary with aggregate morphology. (**A**) We sought to analyse the effects of ABC on the two different morphologies of Aβ(1-40) aggregates. Solution state 1D ^1^H ^15^N-edited HSQC spectra were acquired of 100 μM ABC (blue). Several 20μM aggregation reactions were pooled and concentrated to give 200 μl of either 100 μM fibrillar aggregates (prior pH 11 treatment, black) or non-fibrillar aggregates (no prior pH 11 treatment, red) in an NMR tube. ^15^N labelled ABC was added to give a final concentration of 100 μM, and the samples were incubated for 2 days at 37°C at 700 rpm shaking. 1D ^1^H ^15^N-edited HSQC spectra were taken. The net signal intensity was observed to decrease in the case of type 1 aggregates, and increase in the case of type 2 aggregates, indicating that the interaction between the chaperone and Aβ(1-40) depends on the specific morphology of the aggregates. (**B**) Translational diffusion coefficients can be obtained from the species associated with NMR signals using pulsed field gradient measurements. For ABC alone, a value of 3×10^−7^ cm^2^ s^−1^ was measured, which corresponds to ABC existing predominantly as a 28mer (blue) **(Baldwin, *et al.*, 2011)**. On addition of type 2 Aβ(1-40) aggregates, the diffusion coefficient was unchanged (black). By contrast, on addition of fibrillar type 1 Aβ(1-40) aggregates, the diffusion coefficient was found to increase (individual spectra shown in **Fig. S5B**). This suggests either that the oligomeric state of free ABC chaperones substantially decreases, providing further evidence that ABC can distinguish between morphological forms. (**C**) TEM micrographs of ABC alone reveal discrete particles **(Baldwin, *et al.*, 2011)**. On addition of ABC to either morphology of mature Aβ(1-40) aggregates, no obvious changes in morphology were observed. This is in contrast to the case where ABC was present during initiation of the aggregation reaction (**Fig. 5**).

Further evidence that ABC interacts differently with type 1 and type 2 aggregates comes from solution-state NMR spectroscopy. Using 1D ^15^N edited ^1^H HSQC spectra, we studied the signal intensity of the flexible C-terminus of ABC **(Carver, et al., 1992)**. This segment of ca. 20 residues is both intrinsically disordered and highly dynamic and has been studied in detail previously, providing mechanistic insight into how isolated ABC interconverts **(Baldwin, et al., 2011; Baldwin and Kay, 2012; Baldwin, et al., 2011; Baldwin, et al., 2012)**. Although 1D NMR spectra do not allow for any residue-specific analysis, they can provide both information about molecular interactions, and the translational diffusion coefficient of the species under observation. When incubating ABC with type 1 fibrillar aggregates, the signal intensities of ^15^N-labelled ABC in 1D-HSQC spectra decreased (**Fig. 6A**) and the diffusion coefficient of residual ABC increased (**Fig 6B, S5B**). This indicates that the unbound ^15^N-ABC, which remains visible after a population has bound, was of a lower oligomeric size than in the isolated ensemble. By contrast, incubating ABC with fibrillar type 2 aggregates resulted in an increase in signal intensity (**Fig. 6A**) but no change in the diffusion coefficient (**Fig. 6B**). The signal loss in presence of fibrillar type 1 aggregates can be attributed to a folding of ABC’s otherwise disordered C-terminus upon binding, whereas the intensity gain in presence of disordered type 2 aggregates can be explained by a change in the interconversion rates of ABC, altering the degree of signal broadening due to chemical exchange as has been previously studied in the context of isolated chaperones **(Baldwin, et al., 2011; Baldwin and Kay, 2012; Baldwin, et al., 2012)**. Both findings are consistent with a weak interaction between ABC and Aβ that critically depends on the specific morphology of the Aβ aggregates present.

## Discussion

The assembly of Aβ into amyloid plaques is associated with AD **(Thal, et al., 2015)**, and different morphologies and aggregate sizes have been linked to variations in toxicity **(Bucciantini, et al., 2002; Hochberg, et al., 2014)**. Detailed mechanistic studies have been hampered by the lack of a source of native Aβ that reproducibly forms well-defined morphologies of amyloid. Our protocol effectively solves this problem by providing highly pure Aβ from a recombinant source in relatively high yield.

Working with peptides produced using solid phase synthesis commonly results in batch-to-batch differences, which is likely caused by fluctuations in reaction conditions and the near random occurrence of specific impurities. This makes it a challenging and expensive source for mechanistic studies. Aβ(1-40) produced from our protocol aggregates in a highly reproducible fashion. Moreover, by systematically altering the solution conditions, we can reliably produce amyloid fibrils (prior pH 11 treatment, [Aβ] ≤ 20 μM), which we term here type 1, disordered type 2 non-uniform aggregates (no pH treatment), or a mixture of both (prior pH 11 treatment, [Aβ] > 20 μM). Interestingly, the Aβ-concentration window of 5 – 20 μM, found to form amyloid fibrils only, corresponds to the concentration of insoluble Aβ(1-40) in the superior frontal gyrus and entorhinal cortex of AD patients (**Lue, et al., 1999**).

We demonstrate the importance of this distinction by revealing an unexpected mechanism of action for the chaperone ABC. The links of ABC to AD are well established, **(Ecroyd and Carver, 2009)** but the molecular mechanism behind this role is not well understood. Our data provide insight into this. The surfaces of Aβ fibrils are important for amyloid formation as they provide a template that accelerates aggregate growth, following a kinetic process termed ‘secondary nucleation’ **(Cohen, et al., 2013)**. We can use this principle to explain our results. We suggest that the final morphology of Aβ aggregates reflects a competition between terminal end-growth, that ultimately results in amyloid fibrils (type 1), or a lateral surface mediated assembly process that if dominant, would be expected to lead to predominantly disordered assemblies (type 2). The prior pH 11 treatment and low concentrations favour terminal elongation and amyloid fibrils, whereas higher concentrations together with the lack of pH treatment appear to favour the lateral surface mediated growth and the formation of disordered assemblies, that nevertheless bind the dye ThT, a characteristic of amyloid (**Fig. 7**).

**Fig. 7:**
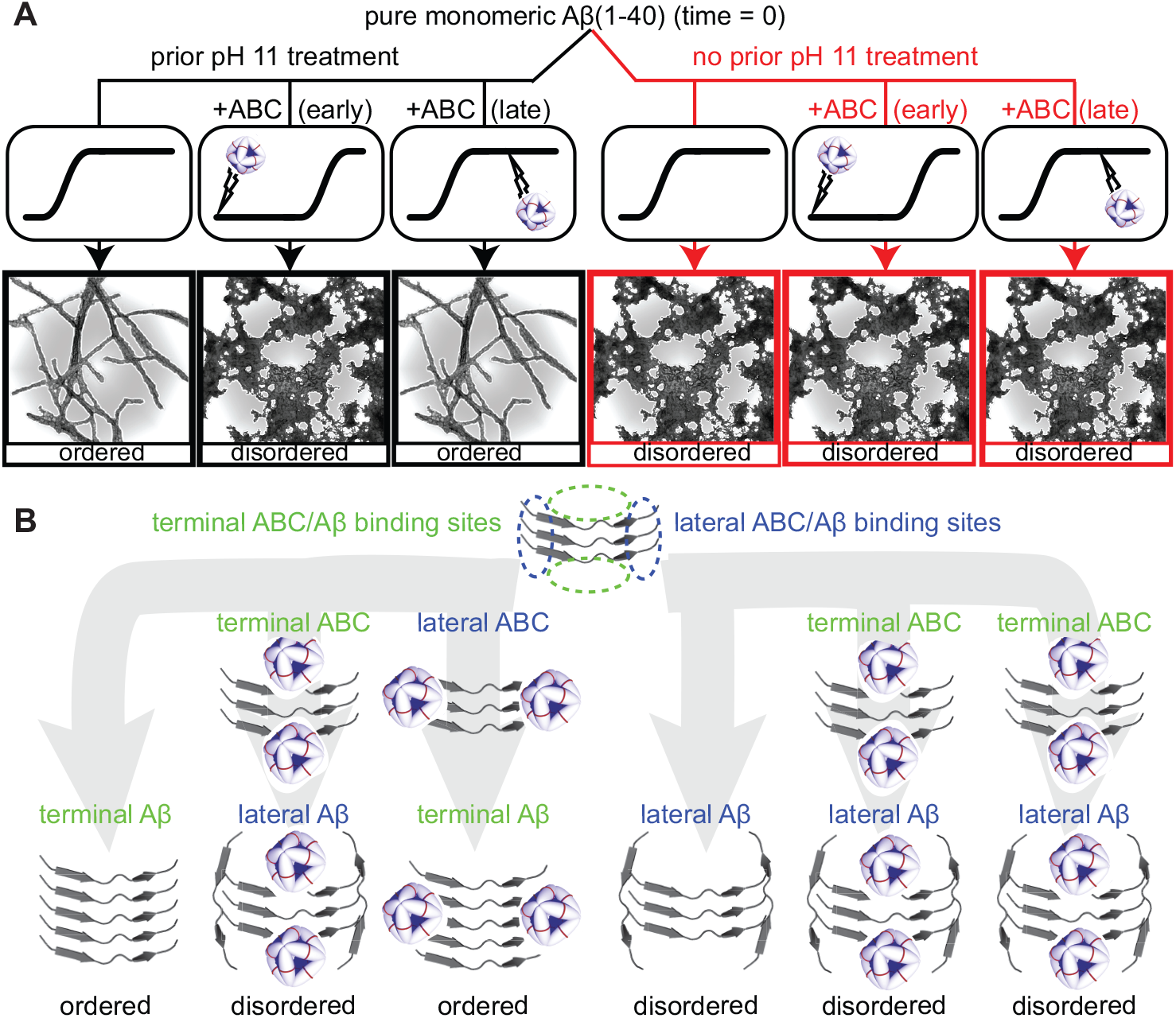
Proposed model for interactions between ABC and Aβ(1-40). Initially, before aggregation, Aβ(1-40) will be predominantly monomeric. When there is a prior treatment at pH 11, fibrillar type 1 amyloid fibrils predominate. When there is no initial treatment, disordered aggregates are observed. We can interpret this by noting that there are at least two possible interaction modes for Aβ(1-40). One involves binding on the terminus of a fibril, leading to ordered growth and amyloid fibrils, and a second involves binding laterally to the surface, likely by a similar process that has been identified as important for secondary nucleation in amyloid formation **(Cohen, *et al.*, 2013)**. The pH treatment affects the relative affinities and timescales of these two modes. We can explain our ABC binding data if we follow the same scheme. Adding ABC early in the aggregation reaction results in disordered growth, whereas adding it to pre-formed aggregates results in no discernable change to the aggregate morphology. We propose that ABC can similarly bind either laterally or terminally to Aβ(1-40) aggregates with comparable affinities. If ABC competes for the terminal-binding site with free Aβ, then growth would be expected to follow the more disordered, lateral aggregation pathway. By contrast, when ABC is added to mature amyloid fibrils, there is a relatively large number of lateral binding sites available, and so binding occurs principally along the length of the fibril **(Shammas, *et al.*, 2011)**.

Similarly to Aβ binding in secondary nucleation, ABC has been shown previously to bind the lateral surfaces of mature amyloid fibrils **(Narayanan, et al., 2006; Shammas, et al., 2011)**. While incubation of ABC with Aβ slows aggregation under all conditions, our results show that mixing ABC and Aβ at the start of the aggregation reaction drastically affects the final morphology of aggregates, with amyloid fibrils being disfavoured and more disordered aggregates becoming prominent. We can rationalise this in terms of the same lateral, and terminal binding as used to describe Aβ amyloid assembly **(Cohen, et al., 2013)** and by extending the observation that there exists competition between Aβ and ABC for the two interfaces **(Narayanan, et al., 2006)**. Our data suggest that ABC effectively slows Aβ binding both surfaces, although the terminal binding appears to be more highly destabilised. On a molecular level, this could be accomplished in principle by ABC either affecting the fold of monomeric Aβ, thereby making it less amenable for elongation, by blocking elongation sites on amyloid fibrils or by promoting lateral assembly of fibrils. It has been previously proposed that ABC interacts with different morphologies with different binding sites **(Mainz, et al., 2015)**, but to our knowledge this has not been shown previously by examining different morphologies of a single aggregating protein. Experimentally distinguishing between these specific effects is a challenging prospect, and will be the subject of further work.

Consistent with ABC being able to distinguish between aggregate morphologies, we see different effects on its C-terminus by solution-state NMR spectroscopy. When we incubate ABC with mature type 1 fibrils, we see a loss of signal intensity, that likely corresponds to a bound population of ABC, and a small reduction in translational diffusion coefficient. These observations suggest that higher molecular weight oligomers have a stronger tendency to bind fibrils, and that the C-terminus folds on binding, leaving a population of smaller oligomers in solution after association. By contrast, signal intensity is observed to rise slightly on incubation with type 2 aggregates, suggesting that the internal subunit exchange dynamics are affected. This indicates a different mode of interaction. It is interesting to note that an ABC signal increase through interaction with one aggregate type and an decrease through interaction with the other aggregate type could result in zero change in ABC signal intensity, if investigated in a mixture of both aggregate types. This has potentially obscured observation of this effect in previous studies. Through preparing well-defined morphologies of Aβ aggregates, as our protocol allows, it becomes possible to quantitatively analyse the interaction of ABC with both morphologies.

Overall, we provide here a new protocol that provides recombinant native Aβ (1-40) peptide in relatively high yield. We show that we have unprecedented control of amyloid morphology using this material. To our knowledge, this is the first time kinetic data of this reproducibility has been routinely produced from the native peptide. We demonstrate the utility of this method by performing a mechanistic study of the interaction of Aβ (1-40) with the chaperone ABC. We show that their interaction affect the resulting morphology of aggregates, with a strong destabilisation of amyloid fibrils in favour of more disordered aggregates, that can be rationalised using a simple binding model (**Fig. 7**). We anticipate that Aβ (1-40) produced using this method will facilitate future structural and mechanistic studies of Aβ peptides and their role in AD.

### Significance

The human small heat shock protein ABC has close links to Alzheimer’s disease. It is up-regulated in neurons and glia adjacent to plaques and is found co-localised in plaques in *ex-vivo* material from AD patients. I*n vitro,* it prevents a wide variety of proteins from forming amyloid, including Aβ peptides as the primary component of AD plaques and it reduces the inherent toxicity of amyloids to cells. Studies of the interaction between ABC and Aβ aggregates have been substantially hindered by challenges associated with producing Aβ-peptides of native sequence which has reproducible aggregation kinetics.

To overcome this, we describe a protocol for producing native Aβ(1-40) recombinantly, in relatively high yields. We demonstrate highly reproducible aggregation kinetics owing to its high purity and that we can produce two different aggregates morphologies under physiologically relevant solution conditions in dependence upon the Aβ concentration and a prior high pH treatment. Type 1 amyloid fibrils can be produced with a prior high pH treatment and an Aβ(1-40) concentration ≤ 20 μM. Type 2 disordered aggregates can be produced in at > 20 μM Aβ(1-40).

We analyse the effects of ABC on the different morphologies of Aβ(1-40) aggregates. When Aβ-aggregation is initiated in presence of ABC incubated at early time points with Aβ, ABC it slows the formation of all aggregate types. Most notably, amyloid fibril formation is inhibited and disordered aggregates are more commonly observed. By contrast, when ABC is added to mature aggregates of either type, it binds but does not affect the morphology. We propose a model to explain these effects based on ABC and Aβ competing for binding to sites on the ends (terminal) or on the sides of fibrils (lateral). We use solution-state NMR spectroscopy to further demonstrate that ABC can distinguish between specific aggregate morphologies. Taken together, these results provide insight into the molecular mechanism that links ABC to AD.

## Experimental Procedures

### Expression and purification

The coding sequence for wild type His-SUMO-Aβ(1-40) was cloned and overexpressed as described in detail in the Supplemental Experimental Procedures. We developed two protocols for purification of Aβ(1-40), **A)**, focusing on denaturing GdnHCl solutions (**Fig. 1B, Fig. 2**) and **B)** using Tris-based buffers (**Fig.1B, Fig. S1**). Both were found to give similar overall yields of up to 4 mg Aβ(1-40) / L *E.coli* culture.. **A)** Cell pellets were suspended in 6 M guanidinium chloride (GdnHCl) pH 8.0 and lysed by microfluidisation. The soluble fraction was loaded onto a Ni^2+^ HisTrap column, washed with 6 M GdnHCl pH 6.0, and eluted with 6 M GdnHCl pH 2.0. This material was loaded onto a Zorbax SB300 C18 column equilibrated in 10 % (vol/vol) acetonitrile (ACN) 10 mM NaP_i_ pH 8 and eluted using an acetonitrile gradient from 10-70% at room temperature. Fractions containing His-SUMO-Aβ were lyophilised and suspended in 50 mM TrisHCl at pH 8. **B)** Cell pellets were suspended in 50 mM TrisHCl 150mM NaCl 1 mM DTT pH 8 and loaded onto a HisTrap column equilibrated in the same buffer. This was eluted using an imidazole gradient to a final concentration of 1 M. Peak fractions were pooled and imidazole removed by sequential rounds of concentration and dilution using centricon (Amicon^®^ Ultra Merck Millipore) columns with a 10 kDa cut-off. **A/B**) Both methods cleave the His-SUMO tag from Aβ(1-40) using the same protocol. The volume is reduced to ca. 200 μl and the tagged Aβ(1-40) is incubated with Ulp1 SUMO-protease (either Invitrogen or produced recombinantly (**Malakhov, et al., 2004; Yates, et al., 2016)**) in a 10:1 His-SUMO-Aβ:Ulp1 concentration ratio at 30 °C for 3 hrs (**A**) or at 4 °C for 2 days (**B**). The efficiency of cleavage was typically 50-80% varying between batches and source of Ulp1. The cleavage products were either diluted to 1 ml using 6 M GdnHCl pH 2.0 (**A**) or loaded directly (**B**) onto a Zorbax SB300 C18 column equilibrated in 10 % (vol/vol) acetonitrile (ACN) 10 mM sodium phosphate pH 8. Highly pure Aβ(1-40) was obtained when eluting with a stepwise acetonitrile (ACN) gradient from 25 - 30 % (vol/vol), incremented in 1% intervals, in 10 mM NaCO_3_ pH 11.0 and a temperature of 40 °C. Uncleaved His-SUMO-Aβ eluted at ACN concentrations higher than 30 % (vol/vol). Pure Aβ fractions were pooled and lyophilised. Poor resolution of His-SUMO-Aβ and Aβ occurred when the column was overloaded. Pooling any mixed fractions, lyophilising, and repeating the column allowed additional recovery of Aβ. Only freshly lyophilised Aβ(1-40) was utilised for aggregation experiments, as even storage in 10 mM NaOH at 4 °C overnight led to non-reproducible aggregation kinetics.

Human αB-crystallin (ABC) was cloned and expressed in M9 minimal medium. ^15^N labelled protein for HSQC spectra (**Fig. 6**) was obtained by providing ^15^N ammonium chloride as the only nitrogen source. ^2^H/^13^CH_3_ Ile δ_1_, Val γ_1_/γ_2_, Leu δ1/δ_2_ labelled ABC was used for binding studies and diffusion analyses (**Fig. 6, S5B**) and was prepared as described previously **(Baldwin, et al., 2011)**, supplementing D_2_O for H_2_O, in presence of ^2^H/^12^C glucose, and adding specific keto-acid precursors 1 h before induction. ABC was purified as described previously by anion exchange chromatography followed by size exclusion chromatography (**Hilton, et al., 2013**).

Synthetic Aβ(1-40) was purchased from AnaSpec Inc. (CA, USA, purity index > 95%) and stored at −20 °C until reconstitution in 10 mM NaOH for aggregation reactions (**Fig. 3**).

### Aggregation kinetics

Two methods were employed to initiate aggregation. Recombinant Aβ(1-40) was first dissolved in either 10mM NaPi 100 mM NaCl pH 7.4 (**Fig. 3,4,5,S3**) or in 50 mM Na-Acetate pH 8.5 (**Fig. S4**). Residual NaCO_3_ from the reverse phase were removed using a PD-10 desalting column (GE Healthcare Life Sciences). Before aggregation, where specified, samples were subjected to a high pH treatment for 30 min at 4 °C with 10 mM NaOH (phosphate) or 10 mM NH_4_OH (acetate). The solution was then taken to pH 7.4 (phosphate) or 8.5 (acetate) with 10 mM HCl.

Aggregation reactions with Aβ(1-40)-concentrations of 5 - 80 μM in absence or presence of 5 μM ABC with a total volume of 150 μl were monitored by light scattering at 340 nm and thioflavin T (ThT)-fluorescence at 485 nm after excitation at 440 nm in sealed Corning 3881 96-well clear bottom polystyrene, nonbinding plates in a FluoStar Omega microplate reader (BMG Labtech, UK) at a temperature of 37 °C with linear shaking at 700 rpm. Changes of light scattering and ThT-fluorescence intensity in absence of protein were negligible. Note that all aggregation traces in this paper (**Fig 4,5,S3,S4**) have scattering and ThT fluorescence normalised to the same scale. This allows to compare all traces directly both within and between all figures. The times required for half completion of a givenaggregation process were obtained by linear regression of the respective data points between ThT fluorescence values of 0.3 and 0.7 (**Fig. 2**). All reactions were performed in triplicate. The typical spread of the data with error bars is shown in **Fig 3, 5**.

### EM

Carbon-coated CF300-CU-50 grids (Electron Microscopy Sciences, Hatfield, USA) were glow-discharged with a Quorum Technologies Q150R ES coating system just before use. 5 μl of solution containing either 5 μM isolated ABC, 20 μM Aβ(1-40) (**Fig 3,4,5,S3,S4**), or 80 μM Aβ(1-40) (**Fig S3**) were deposited on grids and stained with 2 % uranyl acetate as described previously **(Ohi, et al., 2004)**. Images were recorded using a FEI Tecnai 12 transmission electron microscope (TEM) equipped with a lanthanum hexaboride (LaB6) electron source operated at 120 kV and 20 × to 300,000 × magnification, and a bottom-mounted 4 Megapixel Gatan Ultrascan™ 1000 CCD camera.

### NMR

All NMR-spectra were acquired using a triple resonance room temperature probe at a static magnetic field of 14.1 T (Agilent Technologies) at 25 °C in 5 % D_2_O. Protein concentrations were 100 μM for ABC and 100 μM for Aβ(1-40), respectively, with samples prepared through combining and concentrating sets of 20 μM preparations as described (**Fig. 6**). All data sets were zero-filled, phased, and Fourier-transformed with NMRPipe **(Delaglio, et al., 1995)**. Diffusion experiments were conducted as described elsewhere **(Baldwin, et al., 2015)**. In brief, a pulsed field gradient stimulated echo (PFGSE) sequence was used with diffusion delays of both 100 and 200 ms and gradients of a duration of 100 or 200 ms were applied with a strength varying from 4 to 60 G cm^−1^. Both ^1^H, ^13^C, and ^15^N edited experiments were acquired in conjunction with appropriately labelled ABC to ensure that detected NMR signals and corresponding diffusion coefficients originated from the chaperone. Diffusion coefficients were calculated by taking individual intensities and fitting to the Stejskal-Tanner equation **(Stejskal and Tanner, 1965)**, which relates signal intensities to the applied gradients, and the diffusion delay. Raw spectra exemplifying the trends are presented in **Fig. S5B**. The reported diffusion coefficients were obtained by performing ^1^H/^13^C and ^15^N experiments with 1-2 values for the diffusion delays and ≥10 values for the strength of the applied gradients. Uncertainty values were obtained from taking the standard deviation of these independent measurements. This analysis was performed using in-house software programmed in Python, available on request. The self-diffusion of water was also monitored with each measurement to detect any changes in the solution viscosity. No significant variations in viscosity were encountered in this work. For ^1^H experiments, 8k points were typically acquired with a total acquisition time of 0.5 s and 128 scans per transient. For ^13^C and ^15^N edited experiments, 1k points were typically acquired with a total acquisition time of 64 ms and 256 scans per transient.

## Author Contributions

The Aβ(1-40) purification scheme was devised by AJB, and the GdnHCl purification method was developed by HM. Purification conditions were optimised by HM, AvdZ, and DD. The Tris purification method was developed by DD. Aggregation reactions were performed by HM. The manuscript was written by HM and AJB with contributions from all authors.

## Acknowledgements

We are grateful to Dr. Alaji Bah for discussions regarding design of the DNA construct and Dr. Errin Johnson, bioimaging facility, Sir William Dunn School of Pathology, Oxford, for technical assistance. The research was funded by BBSRC grant BB/J014346/01, a David Phillip’s fellowship awarded to AJB. No conflicts of interest were disclosed.

