## Supplemental Information for "αB-crystallin affects the morphology of Aβ(1-40) aggregates"

[1]Physical & Theoretical Chemistry Laboratory (PTCL)
Department of Chemistry
University of Oxford
Oxford OX1 3QZ
UK

### Supplemental Experimental Procedures

*Cloning and overexpression of His-SUMO-Aβ(1-40)*

A DNA construct comprising a His-tag, a SUMO-tag, and the wildtype sequence for Aβ(1-40) was cloned into a pET-28a(+)-plasmid using the NdeI and HindIII cloning sites with coding sequence:

CATATGGGCTCGTCGCATCATCATCATCATCATGGCTCGGGTCTGGTGCCGCGTGGCTCAGCGTCAATGTCGG
ACTCGGAAGTGAATCAAGAAGCGAAACCGGAAGTGAAACCGGAAGTTAAACCGGAAACCCATATTAACCTG
AAAGTTAGTGATGGCAGCTCTGAAATTTTCTTTAAAATTAAGAAAACCACGCCGCTGCGTCGCCTGATGGAAG
CGTTTGCCAAACGTCAGGGCAAAGAAATGGATTCCCTGCGTTTCCTGTATGACGGTATTCGCATCCAGGCGGA
TCAAACGCCGGAAGATCTGGACATGGAAGATAATGACATTATCGAAGCACATCGTGAACAGATCGGCGGTGA
TGCTGAATTTCGCCACGACAGCGGTTACGAAGTCCATCACCAAAAACTGGTGTTTTTCGCCGAAGATGTCGGT
TCAAACAAAGGTGCTATTATTGGTCTGATGGTGGGTGGTGTGGTGAAGCTT

After transformation of the plasmid into *E. coli* Bl21 (DE3) cells and overnight incubation at 37 °C one colony was used to inoculate 50 ml of LB medium supplemented with 50 µg/ml kanamycin and incubated overnight at 37 °C with 200 rpm agitation. Two-litre conical flasks, each with 1 litre LB, were inoculated with 10 ml of overnight culture and induced with 0.75 mM isopropyl β-d-1-thiogalactopyranoside (IPTG) once an OD at 600nm of 0.8 was reached. Expression was allowed at 30 °C for 6 h before cells were harvested by centrifugation (5,000 x g for 10 min at 4 °C). Pellets were stored at -80 °C. For purification, pellets were suspended in 6 M GdnHCl pH 8.0 (denaturing method A Fig 2, methods) or 50 mM TrisHCl 150 mM NaCl pH 8.0 (Tris method B Fig S1, methods) and lysed at 4 °C with several passes through a microfluidizer (M-110L, Microfluidics). Remaining insoluble cell particles were removed by centrifugation (20,000 x g for 30 min at 4 °C) and the soluble fraction was retained.

### Supplemental Figures

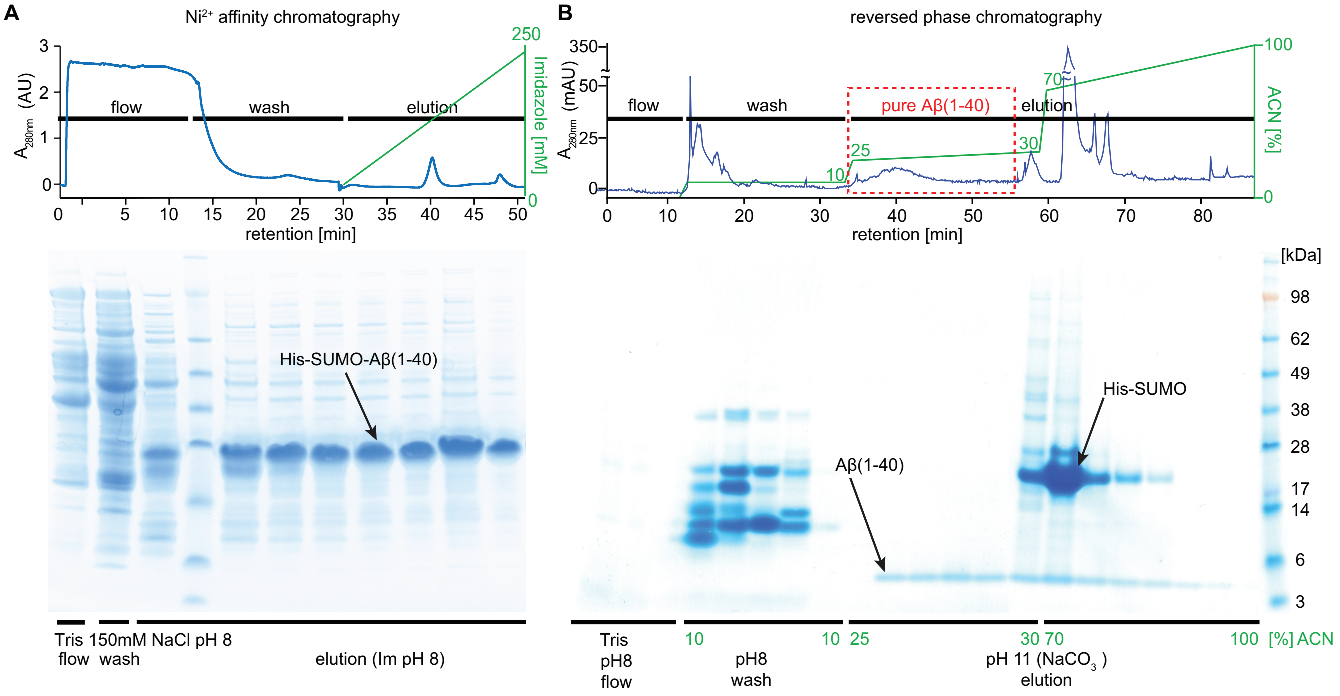


**Fig. S1: An alternative, non-denaturing purification protocol for Aβ(1-40)**. In the course of optimising the purification, we developed two protocols for purification that give similar yields overall. The peptide yields produced from both protocols was found to be identical. The first method uses denaturing buffers (Fig 2, methods). An alternative method in Tris is described here. (**A**) In the Tris method, cells are lysed in 50 mM Tris 100 mM NaCl, pH 8. The cell lysate is loaded on a $Ni^{2+}$ column and washed by adding 50mM imidazole to the running buffer. His-SUMO-Aβ is eluted using an imidazole gradient up to 1 M. The resulting material was concentrated and mixed with Ulp1 for cleavage of the SUMO-Aβ linker as described in the methods. (**B**) The solution containing the digested products is loaded onto a C18 column held at 40 °C and washed with 10 % (vol/vol) acetonitrile 10 mM $NaP_i$ pH 8.0, before elution of pure Aβ(1-40) with a stepwise gradient of acetonitrile to resolve Aβ from uncleaved His-SUMO-Aβ. When the column is overloaded, resolution between these components was poor. Lyophilising unresolved fractions and repeating the reverse phase protocol yielded additional monomeric Aβ.

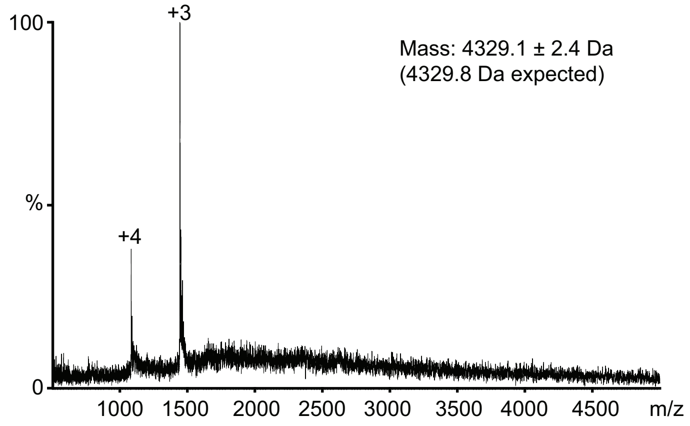


**Fig. S2:** Electrospray ionisation mass spectrometry revealed the purity of Aβ(1-40). In the mass spectrum, only two charge states were clearly visible which together provided an accurate mass reading of 4329.13 ± 2.38 Da. This is in excellent agreement with the expected mass for Aβ(1-40) of 4329.8 Da.

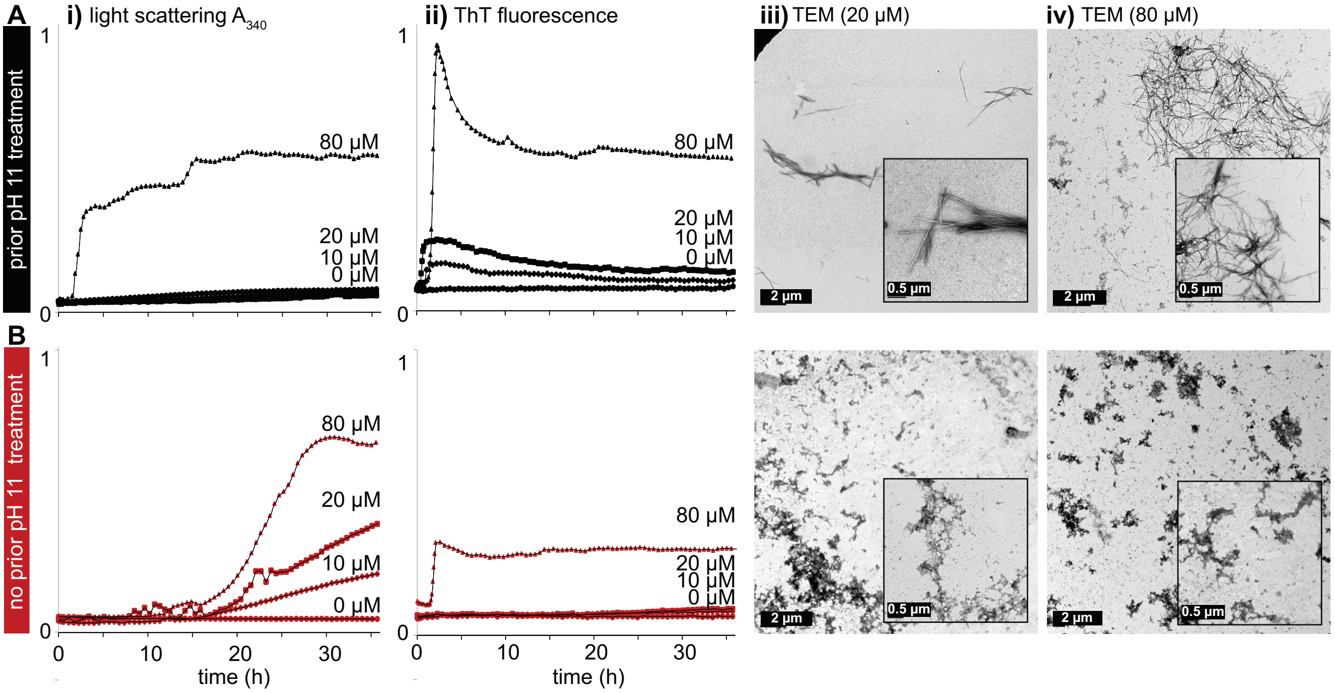


**Fig. S3: Comparison of aggregation at low and high concentrations of Aβ(1-40) in phosphate buffer with and without prior pH 11 treatment.** Light scattering (i), ThT fluorescence (ii), and TEM (iii,iv) were used to follow the aggregation of A→B←(1-40) with (**A**) and without (**B**) prior treatment at pH 11 using 10 mM NaOH for 30 minutes at 4 °C. All aggregation reactions were conducted at 37 °C in 10 mM $NaP_i$, 100 mM NaCl at pH 7.4. The normalisation for the light scattering and ThT fluorescence data are identical for all data shown in this paper (Fig 4,5,S4) to allow quantitative comparisons. At concentrations at or below 20 µM after prior pH 11 treatment, discrete amyloid fibrils (type 1) were observed by TEM. At 80 µM, an appreciable increase in scattering together with an increase in ThT fluorescence was observed. TEM micrographs showed non-fibrillar aggregates in addition to discrete fibrils. To prepare pure amyloid fibrils, the concentration should be kept at or below 20 µM. Error bars have been omitted for clarity. All measurements were conducted in triplicate, and representative variation in kinetic data can be seen in Fig 3 and Fig 5.

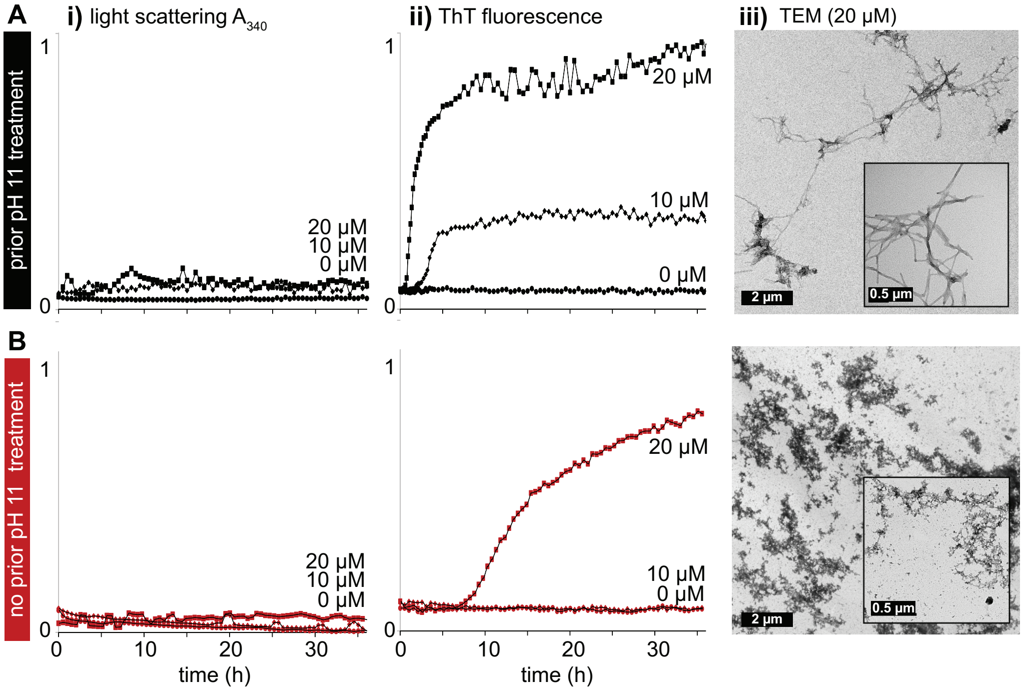


**Fig. S4: The choice of salt used for aggregation reactions can affect the aggregation reaction.** Light scattering (i), ThT fluorescence (ii), and TEM (iii,iv) were used to follow the aggregation of Aβ (1-40) with (**A**) and without (**B**) prior treatment at pH 11 using 10 mM $NH_4OH$ for 30 minutes at 4 °C. All aggregation reactions were conducted at 37 °C in 50 mM Na-Acetate pH 8.5. The normalisation for the light scattering and ThT fluorescence data are identical for all data shown in this paper (Fig 4,5,S3) to allow for quantitative comparisons. The aggregation behaviour in phosphate buffer (Fig 4) and acetate were similar although the duration of the lag times was found to vary. With prior pH 11 treatment, discrete amyloid fibrils were observed, whereas without the treatment, non-uniform non-fibrillar aggregates were obtained. To prepare pure amyloid fibrils, the concentration should be kept at or below 20 μM. Error bars have been omitted for clarity. All measurements were conducted in triplicate and representative variation in kinetic data can be seen in Fig 3 and Fig 5.

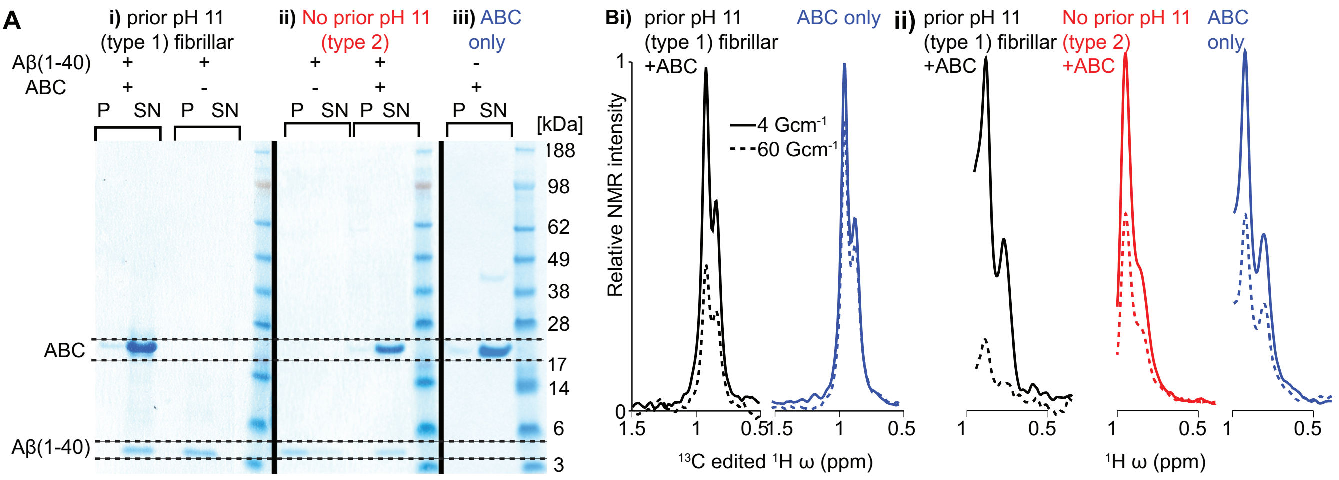


**Fig. S5: Interactions of ABC with Aβ(1-40) aggregates. A)** Differential centrifugation experiments reveal interactions between ABC and Aβ(1-40). Samples of 20 μM Aβ(1-40) and 5 μM ABC were mixed before initiating fibrillation or amorphous aggregation, incubated for 36 hours, and centrifuged at a relatively low centrifugal force of 21,000 x g for 1 h at 4 °C. In absence of ABC, Aβ(1-40) aggregates were found predominantly in the pellet fractions (P), whilst isolated ABC was present in the supernatant fractions (SN). In the mixtures, the majority of Aβ(1-40) was relocated from the pellet to the supernatant fractions, indicating an interaction between both proteins and a decrease in the size of Aβ(1-40)-aggregates. **B)** Selected raw NMR diffusion data that form the basis of the diffusion analysis in Fig. 6. Adding ABC to different morphologies of Aβ(1-40) aggregates results in different diffusion coefficients. $^{13}$C edited $^{1}$H of $^{13}CH_3$ ILV labelled ABC (i) and $^{1}$H (ii) diffusion weighted NMR spectra were acquired in samples containing 100 μM Aβ(1-40) aggregates and 100 μM ABC (**Fig. 6**). When the translational diffusion coefficient of the species under observation increases, the signal attenuation moving from low (solid lines) to high (dotted lines) applied field gradients increases. Once ABC is in the presence of amyloid fibrils, the diffusion coefficient of observed ABC is significantly reduced. By contrast, the translational diffusion of isolated ABC, and ABC in the presence of type 2 aggregates is highly similar. The quantitative diffusion coefficients extracted from spectra acquired with a range of 10 gradient strengths and 2/3 values of Δ are shown in **Fig. 6**. For the data shown, the diffusion delay Δ =200 ms, and the gradient duration δ= 1 ms (i) and δ = 2 ms (ii) was used.
